# Big Data, Small Bias: Harmonizing Diffusion MRI-Based Structural Connectomes to Mitigate Site-Related Bias in Data Integration

**DOI:** 10.1101/2024.10.08.617340

**Authors:** Rui Sherry Shen, Drew Parker, Andrew An Chen, Benjamin E. Yerys, Birkan Tunç, Timothy P.L. Roberts, Russell T. Shinohara, Ragini Verma

## Abstract

Diffusion MRI-based structural connectomes are increasingly used to investigate brain connectivity changes associated with various disorders. However, small sample sizes in individual studies, along with highly heterogeneous disorder-related manifestations, underscore the need to pool datasets across multiple studies to be able to identify coherent and generalizable connectivity patterns linked to these disorders. Yet, combining datasets introduces site-related differences due to variations in scanner hardware or acquisition protocols. These differences highlight the necessity for statistical data harmonization to mitigate site-related effects on structural connectomes while preserving the biological information associated with participant demographics and the disorders. While several paradigms exist for harmonizing normally distributed neuroimaging measures, this paper represents the first effort to establish a harmonization framework specifically tailored for the structural connectome. We conduct a thorough investigation of various statistical harmonization methods, adapting them to accommodate the unique distributional characteristics and graph-based properties of structural connectomes. Through rigorous evaluation, we demonstrate that the generalized linear model with a log-linked gamma model (gamma-GLM) outperforms other approaches in modeling structural connectomes, enabling the effective removal of site-related biases in both edge-based and downstream graph analyses while preserving biological variability. Two real-world applications further highlight the utility of our harmonization framework in addressing challenges in multi-site structural connectome analysis. Specifically, harmonization with gamma-GLM enhances the generalizability of connectome-based machine learning predictors to new datasets and increases statistical power for detecting group-level differences. Our work provides essential guidelines for harmonizing multi-site structural connectomes, paving the way for more robust discoveries through collaborative research in the era of team science and big data.

## 1. Introduction

Diffusion-weighted MRI (dMRI) [1] offers a non-invasive approach for characterizing the brain’s structural connectivity, represented as a structural connectome. Connectomic studies have greatly advanced our understanding of the underlying effects of various brain disorders, including autism spectrum disorder (hereafter, autism) [2, 3], schizophrenia [4], and traumatic brain injury [5, 6]. However, high inter-individual heterogeneity and typically small sample sizes of single-site studies have often led to divergent findings across research efforts [7-11]. In response, the scientific community has embraced big data initiatives by pooling data from multiple clinical sites, with the goal of offering a more comprehensive representation of disorder heterogeneity.

However, substantial site-related variability has been observed in dMRI data collected from different sites, driven by different scanners, acquisition protocols, preprocessing pipelines, and technical differences even with the same manufacturers [12-14]. This site-related variability affects downstream analyses, compromising the comparability of connectome data and obscuring biologically meaningful associations in statistical analyses [15-17]. Various harmonization solutions have been proposed to mitigate site-related biases, targeting the discrepancies at different stages of the data collection and analysis pipelines. For instance, efforts to standardize acquisition protocols and parameters have reduced site-related effects [18-20]. However, this approach cannot resolve site differences retrospectively when data is already collected, and considerable multi-site variations may still arise from residual factors, such as hardware imperfections, operator-specific characteristics, and other uncontrollable variables [18, 21, 22]. Others have attempted to mitigate site-related variations by harmonizing raw dMRI data at the signal level. Mirzaalian et al. [23] introduced a technique that transformed dMRI signals into spherical harmonic space, enabling site-to-site mappings by matching their rotation-invariant spherical harmonic (RISH) features. Subsequently, deep learning methods based on RISH features have also been developed for harmonization of dMRI signals [24, 25]. These RISH-based harmonization approaches have demonstrated effectiveness in mitigating site-related effects in dMRI signals from similar acquisitions. Another approach, proposed by Huynh et al. [26], employs the method of moments (MoM) to directly harmonize dMRI signals without transforming them into other domains, offering greater flexibility for integrating dMRI data with varying numbers of gradient directions.

Despite these advances, most existing approaches for addressing site effects at the dMRI signal level assume that downstream derived measures, such as structural connectomes, will be harmonized accordingly. However, Kurokawa et al. [27], using a traveling-subject design, demonstrated that site-related effects in structural connectomes and their graph metrics can persist even after dMRI signal harmonization. Moreover, different tractography algorithms and connectome construction methods are often adopted in data processing pipelines, particularly for retrospective datasets constrained by acquisition limitations, which can further introduce site-related biases in connectomes. This emphasizes the need for more effective handling of inter-site variability at the connectivity matrix level to adequately mitigate site-related biases in structural connectomes before conducting graph analyses or other investigations in multi-site studies.

Structural connectomes represent complex brain connectivity that is high-dimensional and graph-structured, with connectivity information following a non-normal distribution. As a result, connectome harmonization requires specialized methods to address this complexity. While various statistical models have been proposed for harmonizing neuroimaging data, their applicability to structural connectomes remains largely unexplored. ComBat, originally designed to address ‘batch effects’ in genomic data [28], uses parametric empirical Bayes [29] to adjust for mean and variance of site-related effects while accounting for biological covariates such as age and sex. Fortin et al. [30] extended ComBat to harmonize dMRI-based scalar maps such as fractional anisotropy (FA) and mean diffusivity (MD), and Yu et al. [31] applied ComBat to resting-state functional connectivity. Other widely used harmonization methods include generalized linear model (GLM) [32, 33] and CovBat [34]. Yet, most of these harmonization methods are designed for data that follow a normal distribution. Several studies have applied them to structural connectomes without validating the underlying data assumptions [35], which could lead to incorrect conclusions or unintended consequences in multi-site studies. To date, no harmonization tools have been specifically designed or validated for structural connectomes.

In this work, we address this gap by proposing a harmonization framework for structural connectomes. We propose several statistical harmonization frameworks to accommodate connectome data and conduct a comprehensive evaluation of their effectiveness in eliminating site-related differences while preserving connectivity information and biological associations with age and clinical conditions. To our knowledge, this study is the first effort to propose a tailored harmonization framework specifically for structural connectomes. Furthermore, we present two practical use cases that demonstrate how harmonization approaches can overcome common challenges in multi-site studies. In the first case, we assess the ability of harmonization methods to improve the generalizability of machine learning models to new datasets, particularly in the context of brain age prediction. In the second case, recognizing that most connectome studies include both control and patient groups, we examine the effectiveness of harmonization methods in multi-site data integration to enhance statistical power for detecting clinical group differences. Finally, we provide general guidelines for the harmonization of multi-site structural connectomes, facilitating data integration to yield robust, replicable and generalizable findings.

## 2. Methods

### 2.1. Overview of Datasets

We pooled structural connectome data from six datasets, including the Philadelphia Neurodevelopmental Cohort (PNC) [36], the Center for Autism Research (CAR) [37], Lurie MEGlab [38], and three clinical sites from Autism Brain Imaging Data Exchange II (ABIDEII-NYU, ABIDEII-SDSU, ABIDEII-TCD) [3]. Our study included a total of 1503 participants (890 males, 613 females), comprising 1194 typically developing children (TDC) and 309 children with autism spectrum disorders (ASD). The detailed demographic information and acquisition parameters of each dataset are described in Table 1.

**Table 1.** Demographic information and acquisition parameters for six datasets.

| Dataset | TDC |  |  | ASD |  |  | Scanner | dMRI |  |  |  |  | T1-w MP-RAGE |  |  |  |  |
| --- | --- | --- | --- | --- | --- | --- | --- | --- | --- | --- | --- | --- | --- | --- | --- | --- | --- |
|  | Age range (years) | Sex (M, F) | Sample size | Age range (years) | Sex (M, F) | Sample size |  | b values (s/mm<sup>2</sup>) | Gradient directions | TR (ms) | TE (ms) | Resolution (mm) | TR (ms) | TE (ms) | TI (ms) | Flip angle | Resolution (mm) |
| PNC | 8.17-21.00<br>(15.06±3.33) | M=421<br>F=533 | 954 | - | - | - | Siemen<br>Trio 3T | 0, 1000 | 64 | 8100 | 82 | 1.875×<br>1.875 × 2 | 1810 | 3.5 | 1100 | 9° | 0.9375 ×<br>0.9375 × 1 |
| CAR | 6.26-17.90<br>(11.81±2.85) | M=112<br>F=30 | 142 | 6.37-18.82<br>(12.05±2.73) | M=129<br>F=22 | 151 | Siemens<br>Verio 3T | 0, 1000 | 30 | 11000 | 76 | 2 × 2 × 2 | 1900 | 2.54 | 900 | 9° | 0.8 × 0.8 ×<br>0.9 |
| Lurie<br>MEGlab | 6.17-13.83<br>(9.97±2.04) | M=32<br>F=6 | 38 | 6.17-16.92<br>(10.65±2.18) | M=61<br>F=9 | 70 | Siemens<br>Verio 3T | 0, 1000 | 30 | 11000 | 76.4 | 2 × 2 × 2 | 1900 | 2.87 | 1050 | 9° | 1 × 1 × 1 |
| ABIDEII-NYU | 6.66-12.90<br>(9.55±1.77) | M=19<br>F=0 | 19 | 6.05-17.93<br>(8.33±2.17) | M=35<br>F=4 | 39 | Siemens<br>Allegra | 0, 1000 | 64 | 5200 | 78 | 3 × 3 × 3 | 2530 | 3.25 | 1100 | 7° | 1.3 × 1 ×<br>1.33 |
| ABIDEII-SDSU | 8.10-17.70<br>(13.50±3.05) | M=21<br>F=2 | 23 | 7.40-18.00<br>(12.95±3.35) | M=24<br>F=7 | 31 | GE<br>MR750 | 0, 1000 | 61 | 8500 | 84.9 | 1.875 ×<br>1.875 × 2 | 8.136 | 3.172 | 600 | 8° | 1 × 1 × 1 |
| ABIDEII-TCD | 10.25-20.00<br>(16.18±2.97) | M=18<br>F=0 | 18 | 10.00-19.50<br>(14.47±3.32) | M=18<br>F=0 | 18 | Philips<br>Achieva | 0, 1500 | 61 | 20244 | 79 | 1.94 × 1.94<br>× 2 | 8.4 | 3.9 | 1150 | 8° | 0.9 × 0.9 ×<br>0.9 |

<sup>1</sup>Mean and standard deviation of age are shown in parentheses.
<sup>2</sup>Abbreviations: M, male; F, female, TR, repetition time; TE, echo time; TI, inversion time

In order to evaluate the effectiveness of harmonization methods in different scenarios, we created four data configurations by combining the cohorts in various ways, as detailed below. The overall experiment design and the usage of different data configurations are visualized in Figure 1.

**Figure 1.**
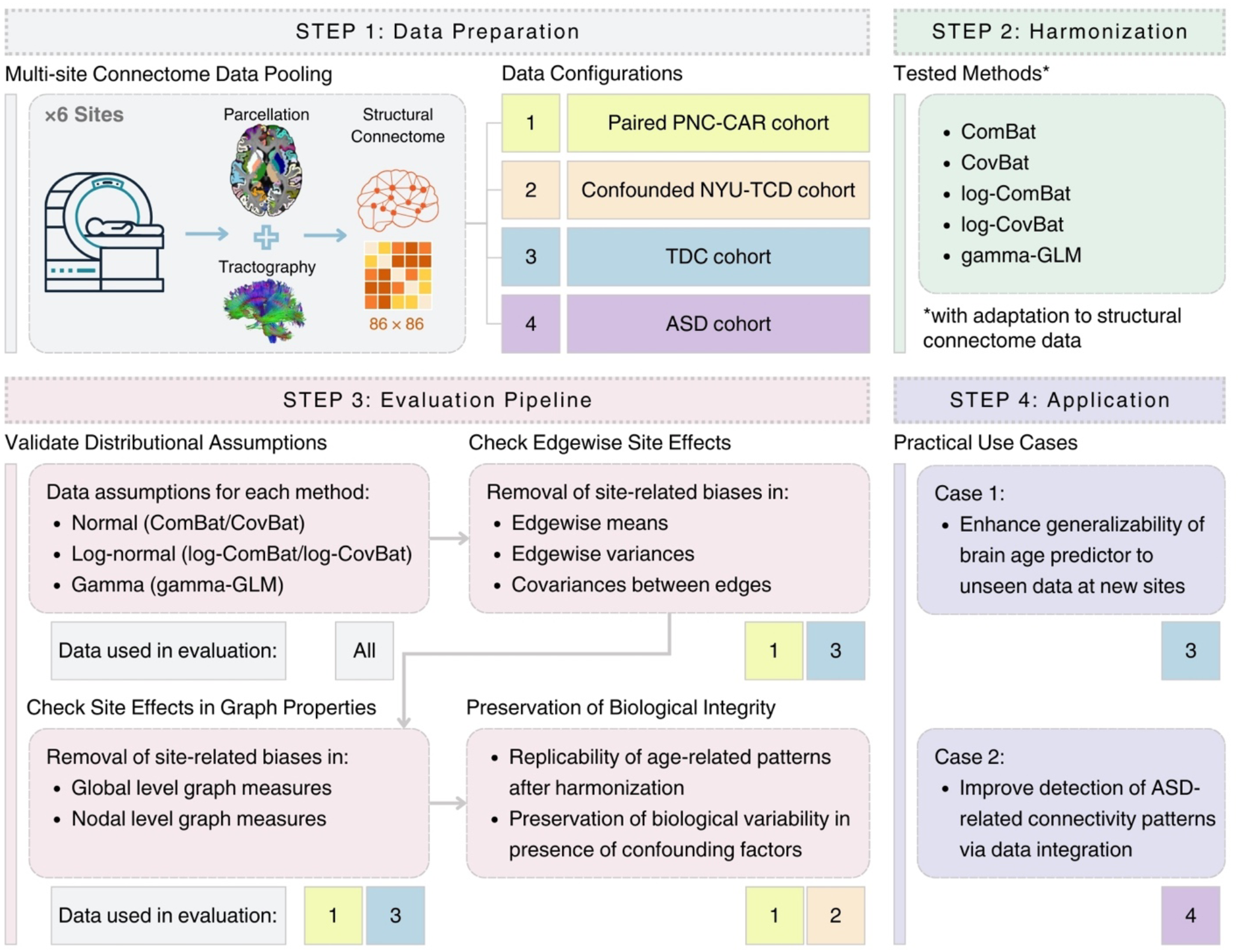
Overview of the harmonization and evaluation framework for structural connectomes. After pooling MRI data from six sites and extracting structural connectomes, we created four distinct data configurations by combining the cohorts in various ways (Step 1). Different harmonization methods were applied to each data configuration for the method comparison (Step 2). We evaluated these harmonization methods from four perspectives: validation of distributional assumptions, removal of edgewise site effects, mitigation of site effects on graph properties, and preservation of biological integrity (Step 3). Finally, we demonstrated the practical utility of structural connectome harmonization methods through two use cases, highlighting their contributions to multi-site studies (Step 4). The data configurations used in each evaluation step are indicated by their corresponding numbers.

#### 2.1.1. Data Configuration 1: Paired PNC-CAR cohort

We carefully selected participants from the TDC group from our two largest datasets, PNC and CAR, ensuring they were closely matched in age and sex. This approach aimed to mimic the traveling-subject research paradigm [23], minimizing confounding effects that could impact the harmonization results. As a result, we included 100 participants (76 males, 24 females) for each site. Participants from the PNC site ranged in age from 8.17 to 18.08 years, with a mean age of 12.31 years and a standard deviation of 2.70 years. Similarly, participants from the CAR site ranged from 8.08 to 17.83 years, with a mean age of 12.27 years and a standard deviation of 2.73 years.

#### 2.1.2. Data Configuration 2: Confounded NYU-TCD cohort

We used all TDC participants from the ABIDEII-NYU and ABIDEII-TCD datasets to create a cohort where age and site were confounded. The ABIDEII-NYU dataset included participants within the youngest age range among the six datasets, spanning 6.66 to 12.90 years, with a mean age of 9.55 years and a standard deviation of 1.77 years. In contrast, the ABIDEII-TCD dataset comprised the oldest participants, ranging from 10.25 to 20.00 years, with a mean of 16.18 years and a standard deviation of 2.97 years. Since both cohorts included only male participants, no sex-related biases were present.

#### 2.1.3. Data Configuration 3: TDC cohort

We pooled all TDC data from our datasets, resulting in a total of 1194 participants (623 males, 571 females). The age of participants ranged from 6.17 to 21.00 years, with a mean age of 14.41 years and a standard deviation of 3.53 years. This cohort represented a more generic case in which age and sex were unbalanced with respect to site, potentially introducing statistical confounding.

#### 2.1.4. Data Configuration 4: ASD cohort

All datasets, except for PNC, included neuroimaging data from children with ASD. We pooled the ASD data from these five datasets to create a cohort of 309 ASD participants (267 males and 42 females) with aged from 6.05 to 19.50 years, with a mean age of 11.49 years and a standard deviation of 3.05 years. The structural connectomes in the ASD cohort were harmonized by adjusting for site effects estimated from the corresponding TDC cohort.

### 2.2. Data Processing

#### 2.2.1. Pre-processing and Quality Control

We conducted a series of preprocessing steps for dMRI scans, including denoising using joint local principal component analysis (PCA) [39], correction for eddy currents and movements with FSL EDDY [40], and skull stripping with BET2 [41]. We preprocessed the high-resolution T1-w images with FreeSurfer recon-all pipeline (http://surfer.nmr.mgh.harvard.edu) [42], followed by registration to the dMRI data of each subject using restricted deformable SyN algorithm in ANTs [43]. Constrained spherical deconvolution (CSD) of dMRI was then conducted using MRtrix3 [44], with the fiber response function estimated using D’Hollander’s algorithm [45, 46] and the fiber orientation distribution (FOD) fitted using the method proposed by Jeurissen and others [47]. We performed probabilistic fiber tracking with a step size of 1 mm and an angle threshold of 60 degree utilizing the iFOD2 algorithm [48] and anatomically constrained tractography (ACT) [48], with 250 seeds placed at random inside each voxel of the grey-matter white-matter interface (GMWMI). We conducted manual quality control to exclude data with poor acquisition quality or failed registration.

#### 2.2.2. Construction of Structural Connectomes

We parcellated the brain into 86 regions of interest (ROIs), including 68 cortical regions from the Desikan-Killiany atlas [49] and 18 subcortical regions from FreeSurfer’s atlas [50]. We calculated the number of streamlines intersecting each pairwise combination of these ROIs, yielding an 86×86 connectivity matrix. Post-processing of structural connectomes involved removing self-connections and symmetrizing the connectivity matrix. Additionally, we normalized the structural connectomes by dividing edge weights by each subject’s total GMWMI volume. Since structural connectomes derived from probabilistic tractography may include numerous spurious connections that could affect downstream analysis, we applied consistency-based thresholding [51] to filter out spurious connections and retained only the most consistent ones across the population. For our primary analyses, we used a density threshold of 40%, while analysis of different consistency levels was also provided in Supplementary Materials.

### 2.3. Harmonization Methods

Based on different distributional assumptions, we investigated five statistical approaches for structural connectome harmonization. Firstly, we tested two widely used harmonization tools, ComBat and CovBat, both of which assumed normal distribution of the data. We then modified the two approaches and introduced log-ComBat and log-CovBat harmonization, which utilized a log-normal data assumption. Finally, we developed a generalized linear model with gamma distributed variables (gamma-GLM) for connectome harmonization, adopting a gamma distribution to fit the connectome data. Table 2 compares these harmonization methods in terms of their capabilities, as also demonstrated in previous literature [28, 34, 52]. All harmonization methods were performed with sex, age, and age^2^ included as covariates.

**Table 2.** Comparison of mainstream harmonization methodologies.

| Harmonization method | Addresses mean site effects | Addresses variance site effects | Addresses covariance site effects | Address skewness | Assure non-negative edge values | Assumption for data distribution |
| --- | --- | --- | --- | --- | --- | --- |
| ComBat | Yes | Yes |  |  |  | Normal |
| CovBat | Yes | Yes | Yes |  |  | Normal |
| log-ComBat | Yes | Yes |  | Yes | Yes | Log-normal |
| log-CovBat | Yes | Yes | Yes | Yes | Yes | Log-normal |
| gamma-GLM | Yes |  |  | Yes | Yes | Gamma |

For each method, we assumed that structural connectomes have been collected from *M* imaging sites, where *m* = 1, 2, …, *M* denotes the index of the site, and each site comprised *N*_*m*_ samples, with *n* = 1, 2, …, *N*_*m*_ representing the index of the subject. *Y*_*mne*_ represents the observed edge value before harmonization at a given edge index *e* ∈ *E*(*G*) within the structural connectome of subject *n* from site *m*. In the following sections, we provided detailed descriptions of each technique for structural connectome harmonization.

#### 2.3.1. Applying ComBat to Connectomes

The ComBat model was originally developed by Johnson et al. [28] to mitigate batch effects when integrating multiple microarray datasets for gene expression analysis. We extended ComBat to harmonize structural connectomes by formulating each edge value as follows:

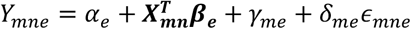

In this model, *α*_*e*_ represents intercept of the connectivity value at edge *e*, the vector ***X***_***mn***_ incorporates covariates of interest, such as age and sex, with ***β***_***e***_ representing the vector of regression coefficients corresponding to ***X***_***mn***_ at edge *e. γ*_*me*_ and *δ*_*me*_ denote the mean and variance of the site effect for site *m* at edge *e*. ComBat assumes that the error term *ϵ*_*mne*_ follows an independent normal distribution 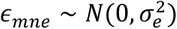. It first estimates 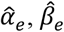 using ordinary least-squares by constraining 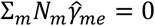 for all edges. To estimate empirical statistical distributions of *γ*_*me*_ and *δ*_*me*_, ComBat employs an empirical Bayes approach with the assumption that all edges share a common yet site-specific prior distribution, with hyperparameters estimated empirically using the method of moments. This allows information from all edges to be leveraged to infer the statistical properties of the site effect. Specifically, *γ*_*me*_ is assumed to have an independent normal prior distribution, and *δ*_*me*_ follows an independent inverse gamma prior distribution. The estimates 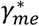 and 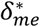 are obtained by computing the conditional posterior means. Finally, ComBat adjusts for the estimated site effects to obtain the harmonized edge:

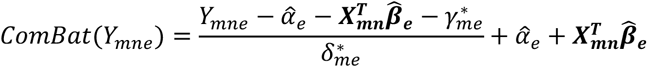

ComBat harmonization may introduce spurious connections for those originally with 0 edge values (indicating no connection between two nodes), altering the physical interpretation of the connectivity. To correct this, we followed the approach of Onicas et al. [35] by reassigning zeros to those connections. Specifically, for each participant, we apply a binary connectivity matrix from the pre-harmonized data to mask the harmonized structural connectomes.

#### 2.3.2. Applying CovBat to Connectomes

ComBat treats each edge in the structural connectomes independently, without accounting for the covariance between edges. However, site effects might be correlated edgewise, exhibiting different covariances across sites. To address these potential covariance effects, Chen et al. [34] adapted the ComBat framework into CovBat. CovBat posits that the error vector ***ϵ***_***mn***_ = (*ϵ*_*mn*1_, *ϵ*_*mn*2_, …, *ϵ*_*mne*_, …, *ϵ*_*mnE*_)^*T*^ ∼ *N*(**0, Σ**_***m***_), where *E* denotes total number of edges. CovBat first applies ComBat to remove the site-specific mean and variance, yielding ComBat-adjusted residuals:

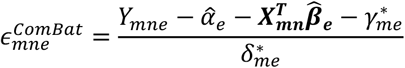

It then performs principal component analysis (PCA) on the residuals. These residuals can be projected onto the principal components, expressed as:

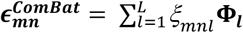

Here, **Φ**_***l***_ represent the eigenvectors of the covariance of the residual data, and 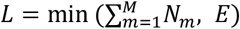 denotes the rank. *ξ*_*mnl*_ indicates the principal component scores. Assuming that the covariance site effect is captured within these principal component scores, they can be harmonized to mitigate the covariance site effects. Suppose:

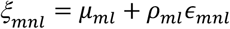

Where *μ*_*ml*_ and *ρ*_*ml*_ are location and scale shifts on principal component scores for site *m* at principal component *l*. The CovBat adjusts these site-related shifts in principal component scores:

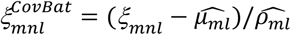

The covariance-adjusted residuals can be obtained by projecting the adjusted principal component scores back into the original space, with a hyperparameter *K* controlling the desired portion of principal component scores to be adjusted:

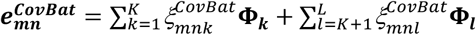

In this study, the hyperparameter *K* was selected such that the principal components explain 95% of the variation. The final harmonized edge values can be represented by replacing the residuals:

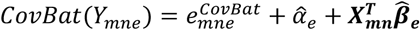

Similar to the post-processing step in ComBat, we reassigned zero values to harmonized connections that originally had zero edge values before harmonization.

#### 2.3.3. Adapting log-ComBat for Connectomes

The ComBat algorithm relies on the assumption of data conforming to a normal distribution. However, this premise may not hold true for structural connectome data, where edge values frequently demonstrate notable skewness. In biomedical research, the log-scaling technique is commonly employed to transform skewed data to approximate a normal distribution [53-55]. The log-scaling approach assumes that edgewise connectome strength follows a log-normal distribution or closely approximate it. Consequently, the ComBat algorithm can be reformulated as follows:

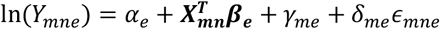

and the harmonized edge values can be obtained from the inverse of a logarithm:

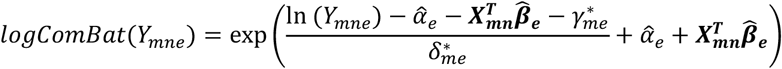

log-ComBat ensures that harmonized edges remain non-negative. To handle numerical errors with the logarithm when original edge values *Y*_*mne*_ are zero, we added a small positive constant *c* to *Y*_*mne*_. Connections with zero edge values prior to harmonization will be reassigned to zeros to ensure no spurious connections are introduced by the harmonization process.

#### 2.3.4. Developing log-CovBat for Connectomes

We also developed log-CovBat, a modification of the CovBat algorithm that incorporates log-scaling to address the skewness of edge weights in structural connectomes. We first applied a log-transformation to all edge values, then performed CovBat on the log-transformed edges. The final harmonized edge values were obtained by applying the inverse logarithm (exponentiation), as formulated below:

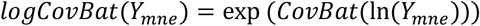

log-CovBat ensures that edges remain non-negative after harmonization. Similarly, we add a small positive constant *c* to original edge value *Y*_*mne*_ during log-CovBat harmonization and restore connections that originally had zero edge values back to zeros.

#### 2.3.5. Developing gamma-GLM for Connectomes

The generalized linear model (GLM) provides an alternative for regression when the assumptions of normality and homoskedasticity are not met [56]. By employing a gamma distribution, GLM offers greater flexibility in modeling a wide range of data shapes, particularly well-suited for right-skewed data. The edge values for each connection are assumed to follow a gamma distribution, *Y*_*mne*_ ∼ Γ(*K*_*e*_, *θ*_*e*_), where *K*_*e*_ and *θ*_*e*_ denote the shape and scale parameters, respectively. The expected value of *Y*_*mne*_ is given by:

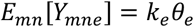

The parameters *K*_*e*_ and *θ*_*e*_ can be fitted using GLM regression. We modeled the effects of site and other covariates on the expected structural connectomes using a log-link function to ensure positive edge values after harmonization, which can be expressed as:

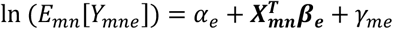

The mean site effects 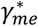 for site *m* at edge *e* can be then estimated. We regressed out the estimated mean site effects 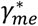 from the edge values *Y* and applied the inverse logarithm transformation (exponentiation):

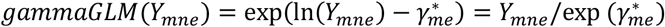

The gamma-GLM with a log-link function ensures that the harmonized edge values remain non-negative. As gamma fitting also requires positive edge values, we add a small positive constant *c* to the original edge value *Y*_*mne*_ and restore connections with zero edge values back to zeros after harmonization.

### 2.4. Validation of Distributional Assumptions on Structural Connectomes

The harmonization methods discussed above are based on three different distributional assumptions for the structural connectome data: the normal distribution (as required by ComBat and CovBat), the log-normal distribution (for log-ComBat and log-CovBat), or the gamma distribution (for gamma-GLM). We conducted goodness-of-fit tests on the structural connectome data to assess the validity of these data assumptions. Specifically, we fitted each statistical distribution to the observed structural connectomes at each site. Kolmogorov–Smirnov (KS) test was used to quantify the discrepancy between the observed data distribution and the fitted distributions.

### 2.5. Evaluation of Harmonization Methods

#### 2.5.1. On Removing Site Effects in Edgewise Connectome Analysis

We first examined whether each harmonization method successfully eliminated site-related biases in the mean, variance, and covariance of edgewise connectome strength. This evaluation utilized the paired PNC-CAR cohort (Data Configuration 1) and the multi-site TDC cohort (Data Configuration 3). Prior to edgewise evaluation, covariates for sex, age, and age^2^ were regressed out. To assess mean site effects, we performed a non-parametric Kruskal-Wallis test to identify edgewise group differences across sites. Variance site effects were evaluated using the Brown-Forsythe test on the residuals of edgewise strength. Brown-Forsythe test is a non-parametric equivalent of Levene’s test to determine whether variances were equal across sites. For site-specific covariance effects, empirical covariance matrices were computed for each site. The Frobenius distance between within-site covariance matrices was then calculated for each site pair, both before and after harmonization. Box-M test was used to determine whether the site-specific covariance matrices were statistically equivalent.

#### 2.5.2. On Removing Site Effects in Derived Graph Topological Measures

Structural connectomes represent brain connectivity as graphs, therefore, effective harmonization approaches should address not only edgewise site effects but also site-related variations in graph properties, which are typically assessed using graph topological measures [57]. We computed graph topological measures at two levels: node-level features (node strength, betweenness centrality, local efficiency, clustering coefficient) and global graph measures (global strength, intra- and inter-hemispheric strength, characteristic path length, global efficiency, modularity). Site-related differences in these measures were analyzed before and after applying harmonization methods using a two-sample t-test for the paired PNC-CAR cohort (Data Configuration 3) and a one-way ANOVA test for the multi-site TDC cohort (Data Configuration 3), after controlling for sex, age, and age^2^. Effect sizes for site differences were calculated using Cohen’s d for comparisons between two sites and Cohen’s f when involving more than two sites.

### 2.6. Evaluation of Biological Variability Preservation

Ideally, the harmonization method should remove non-biological variability introduced by different sites and scanners while preserving the statistical power to detect biologically meaningful associations. To validate this, we calculated the Spearman correlation coefficients between edgewise connectivity strength and participants’ chronological age in the paired PNC-CAR cohort (Data Configuration 1), controlling for sex as a covariate. We expected that the harmonized structural connectomes would retain the age-connectivity relationships observed in the pre-harmonization data for each site. The replicability of age-related patterns was quantified by the true positive rate (sensitivity) of connections showing significant age correlations in the harmonized connectomes compared to the pre-harmonization data. Additionally, we ranked the connections by the absolute values of their age correlations and computed the overlap between the top *K* correlations in the harmonized and pre-harmonization data within each site. We visualized the results using concordance at the top (CAT) curves [58] by varying across all *K* values. A CAT curve closer to one indicates better overlap between the two sets of correlation relationships.

### 2.7. Evaluation of Harmonization with Confounding Factors

Confounding between age and site presents a challenge for harmonization, as removing site-related variation can inadvertently change age-related variation if not handled carefully. To evaluate how different harmonization methods handle this issue, we conducted a Spearman correlation analysis to examine the relationship between edgewise connections and participants’ chronological age within each site using the confounded NYU-TCD cohort (Data Configuration 2). Similar to the previous analysis, CAT plots were used to visualize the replicability of these age associations before and after applying various harmonization methods, while the true positive rate quantified the replicability of connections with significant age correlations. Additionally, we computed the Frobenius distance between the edgewise age correlation matrices of the two sites. We expected this distance would remain stable before and after harmonization, indicating that age-related differences were preserved between sites. Finally, we jointly tested the replicability of age associations after combining data from both sites post-harmonization. We also included a scenario where the two sites were combined without proper harmonization to emphasize the critical role of harmonization in addressing confounding factors during data integration.

### 2.8. Practical Use Cases for Multi-site Connectome Harmonization

To further demonstrate the importance of harmonization in multi-site structural connectome analysis, we evaluated the effectiveness of harmonization frameworks through two practical use cases: 1) improving the generalizability of prediction models to new datasets; we used brain age prediction as an example; and 2) enhancing statistical power for clinical studies through data integration; here we focused on detecting group differences and clinical associations in ASD.

#### 2.8.1. Brain age prediction on new datasets

Brain development is accompanied by structural changes in brain organizations, and the gap between predicted brain age from neuroimaging data and chronological age has been linked to various health conditions [59-63]. While many researchers have used machine learning to model neurotypical developmental trajectories to predict brain age, these predictors often struggle with generalizability across datasets due to site and scanner variability. We used brain age prediction as a case study to assess whether harmonization methods could eliminate site-related effects while preserving age-related biological variability in structural connectomes, thereby improving the prediction performance of machine learning models on new datasets. This evaluation was conducted on the TDC cohort (Data Configuration 3).

We first trained a support vector regression (SVR) model using structural connectomes from the PNC dataset to predict brain age. Features were extracted from the upper triangular portion of the connectivity matrices, with PCA applied for dimensionality reduction. The PNC dataset was randomly split 50:50 into a training set and a testing set. To optimize the model, we applied 5-fold cross-validation on the training set to tune hyperparameters. The final SVR model was trained on all subjects in the training set using the optimal hyperparameters. Model performance was evaluated using the testing set from the PNC dataset. To assess the model’s generalizability, we tested it on unseen data from the remaining sites, evaluating prediction performance through root mean square error (RMSE), mean absolute error (MAE), and Pearson correlation coefficient (R).

#### 2.8.2. Enhancing statistical power for ASD studies

Previous literature has identified several structural connectome patterns associated with ASD [2, 64-67]. However, most studies were conducted at single sites and often had small sample sizes, which limited their statistical power and ability to draw generalizable conclusions. Pooling data from multiple sources can introduce site-related biases that may obscure the detection of clinical associations. Therefore, harmonizing structural connectomes is essential for uncovering reliable ASD-related characteristics in multi-site studies.

To validate the effectiveness of harmonization algorithms, we applied site-specific coefficients, estimated from the TDC cohort (Data Configuration 3), to harmonize the ASD cohort (Data Configuration 4). Given the pronounced sex differences in autistic children and the small sample size of autistic females, we excluded female subjects from this analysis to minimize potential sex bias. We then calculated global graph topological measures of structural connectomes before and after harmonization, assessing group-level differences between the ASD and TDC groups, as well as group-by-age interactions in these measures. Additionally, we conducted the same group-level analyses on unharmonized data, with the site covariate regressed out at the level of graph measures and compared the results with those from the harmonized structural connectomes. We further performed Spearman correlation analysis between graph measures and ASD and developmental clinical assessments, including the Autism Diagnostic Observation Schedule (ADOS) calibrated severity scores, Social Responsiveness Scale (SRS-2), and Vineland Adaptive Behavior Scales (VABS), while controlling for age. Multiple comparisons were corrected using the False Discovery Rate (FDR). As the ComBat and CovBat methods may produce negative edge values, which lack physical relevance when computing graph topological measures in clinical contexts, we excluded these two methods from this analysis.

## 3. Results

### 3.1. Distributional Properties of Connectivity Strength in Connectomes

Each edge of the pre-harmonized structural connectomes was examined to determine whether it followed one of three hypothesized probability distributions: normal, log-normal, or gamma. Figure 2A shows the edgewise KS distances between the observed distributions and three fitted hypothesized distributions in the PNC dataset, the largest site for the TDC group. A larger KS distance indicates a greater deviation of the hypothesized distribution from the observed data. We showed that the gamma distribution demonstrated the best fit, with significantly lower KS distances compared to the other two hypothesized distributions (p<0.0001, paired t-test), followed by the log-normal distribution. Figure 2B illustrates the heatmaps of KS distances at each edge, showing only the edges with significant differences from the observed distributions (p<0.05, FDR-adjusted). Notably, the normality data assumptions underlying ComBat and CovBat were not met at most edges. Among the top 40% of highly consistent edges (1480 edges in total), 97% did not follow a normal distribution, 41% showed significant differences from the log-normal distribution, and only 5% were distributed significantly different from the gamma distribution fitting.

**Figure 2.**
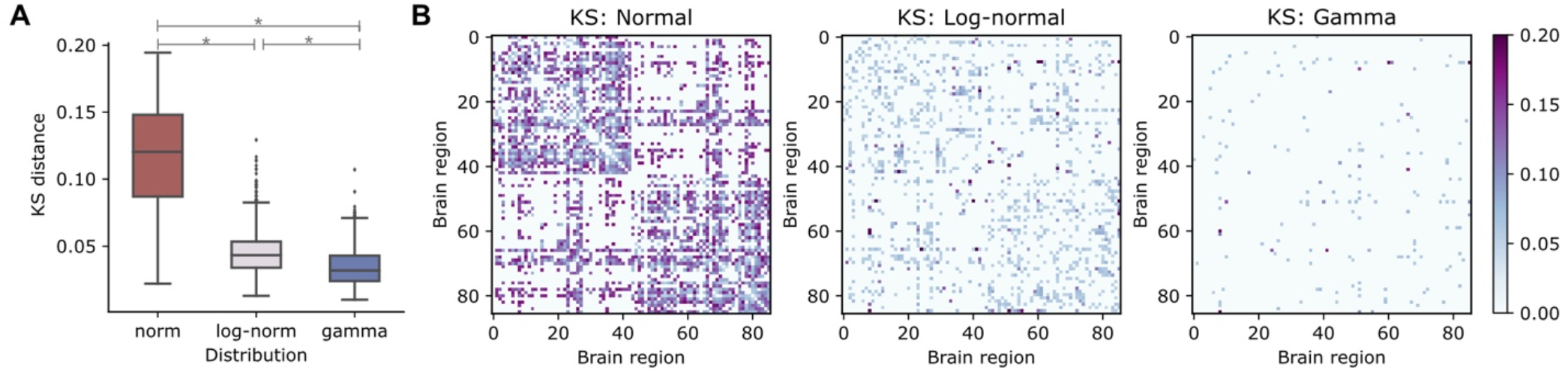
Goodness of fit test for PNC. A) Edgewise KS distances between the observed and three hypothesized distributions (normal, log-normal, and gamma) of connectivity strength in PNC. Asterisks denote significant differences between those hypothesized distributions (p < 0.0001, paired t-test). The gamma distribution provided a significantly better fit than the other two hypothesized models. B) Heatmaps of edgewise KS distances, showing only the edges with significant discrepancy from the observed distributions in pre-harmonized structural connectomes (p < 0.05, FDR-adjusted).

Similar distributional properties of structural connectomes were also observed in the TDC group across the other five sites and held true for the ASD group as well, as shown in Figure S1. Structural connectome data were more likely to follow a gamma distribution, regardless of site or diagnostic group. We also evaluated the goodness of fit by varying different consistency threshold levels, as shown in Figure S2. For highly consistent edges, the gamma distribution provided a significantly better fit than the log-normal distribution (p < 0.0001, paired t-test). For less consistent edges (likely representing spurious connections), the fits of the log-normal and gamma distributions were comparable (p > 0.05, paired t-test). The normal distribution, however, resulted in significantly poorer fits compared to the other two distributions, regardless of the consistency level.

### 3.2. Edgewise Evaluation of Site Effects on Structural Connectomes

We utilized the paired PNC-CAR cohort (Data Configuration 1) to assess the site effects on edgewise strength. Figure 3A displays the MA-plot [68] of site differences between PNC and CAR. The MA-plot is a common method for visualizing differences in measurements taken from two settings, where the edgewise mean connectivity strength was calculated for each site, and the between-site differences in log-transformed means were displayed as a function of the average log-transformed site means. If the structural connectome were free of edgewise site-related effects, the scatterplot would align along the horizontal line at zero, as indicated by the dashed lines. We observed striking site differences in edgewise strength between two sites before harmonization. These site differences were largely reduced after applying any of those hypothesized harmonization approaches.

**Figure 3.**
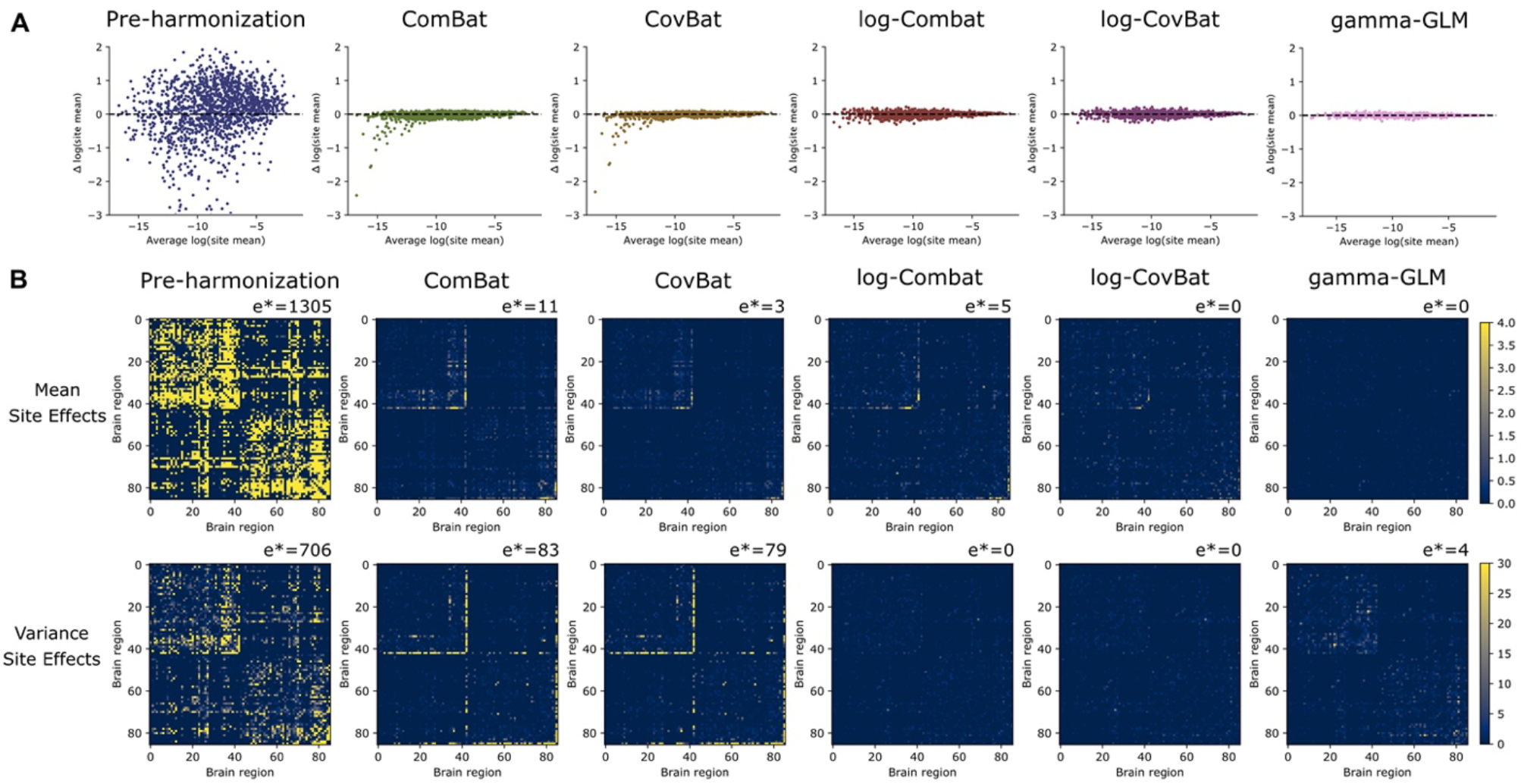
Site-related effects on mean and variance of edgewise strength in the paired PNC-CAR cohort (Data Configuration 1). A) MA-plots for visualization of site differences between PNC and CAR with paired subjects. The x-axis represents the averaged log-transformed means across sites and the y-axis represents between-site differences in log-transformed means. The horizontal line at zero indicates no site-related effects. B) Edgewise site effects on mean (first row) and variance (second row) connectivity strength, showing the Kruskal-Wallis H statistics for mean site effects and F* statistics from the Brown-Forsythe test for variance site effects. The number of edges with significant site effects was noted by e* in the top leB corner of each plot.

Figure 3B illustrates site-related biases in the mean and variance of edgewise connectivity strength, showing Kruskal-Wallis H statistics for mean site effects (first row) and F^*^ statistics from the Brown-Forsythe test for variance site effects (second row). The log-CovBat and gamma-GLM approaches demonstrated favorable performance in eliminating mean site effects, while log-ComBat and log-CovBat were particularly effective in harmonizing variance site effects. Specifically, among the 1480 edges obtained after consistency thresholding, 1035 (70%) edges initially showed significant differences in site means before harmonization (p<0.05, Kruskal-Wallis test). After ComBat harmonization, the number of edges with significant mean site differences dropped to just 11 edges (0.74%), and logComBat reduced the number to 5 edges (0.34%). After CovBat harmonization, only 3 edges (0.20%) showed significant site differences in means, and no significant site effects were observed with log-CovBat and gamma-GLM harmonization. For variance site effects, 706 edges (4.8%) exhibited significant site differences in pre-harmonization data (p<0.05, Brown-Forsythe test). After harmonization, 83 edges (5.6%) still showed significant site effects with ComBat, 79 edges (5.3%) remained significant with CovBat, and 4 edges (0.27%) with gamma-GLM. With log-ComBat and log-CovBat harmonization, all connections showed no statistically significant site-related effects on variance.

Table 3 shows the site-specific covariance effects on edgewise strength before and after harmonization, quantified by the Frobenius distance between covariance matrices of two sites. We regressed out the demographic factors, including sex, age, and age^2^, jointly across both sites from edgewise strength using a linear model. The log-CovBat harmonization approach most effectively mitigated site effects on covariance in the paired PNC-CAR cohort (Data Configuration 1), followed by log-ComBat and gamma-GLM. However, results from the Box’s M test indicated that covariance site effects on structural connectomes were not significant across all tested cases, including the pre-harmonization data.

**Table 3.** Frobenius distances between site-specific covariance matrices.

| Pairwise Site differences | Pre-harmonization | ComBat | CovBat | log-ComBat | log-CovBat | gamma-GLM |
| --- | --- | --- | --- | --- | --- | --- |
| Paired PNC-CAR | 0.1022 | 0.0924 | 0.0912 | 0.0728 | 0.0726 | 0.0786 |

### 3.3. Evaluation of Site Effects on Derived Graph Topological Measures

We examined the derived graph topological measures of structural connectomes to determine whether harmonizing edgewise connectivity strength can lead to harmonized downstream outcomes. Figure 4 illustrates the Cohen’s d effect sizes of site differences across six global graph topological measures, with asterisks denoting significant site effects. All six global measures showed significant site effects before harmonization (p<0.05, two-sample t-test), with three strength-based graph measures exhibiting large effect sizes (Cohen’s d > 1.0), characteristic path length displaying a moderate effect size (Cohen’s d=0.68), and global efficiency showing a small effect size (Cohen’s d=0.30). After ComBat harmonization of edgewise connections, site effects on characteristic path length and global efficiency remained significant (p<0.05, two-sample t-test), though their effect sizes were reduced to small (Cohen’s d=0.20 for characteristic path length, Cohen’s d=0.27 for global efficiency). Among CovBat, log-ComBat, log-CovBat, and gamma-GLM, all of which addressed site effects on each global measure, leaving no measure with significant site-related differences. The gamma-GLM showed the best overall harmonization performance, yielding the smallest effect sizes across most measures.

**Figure 4.**
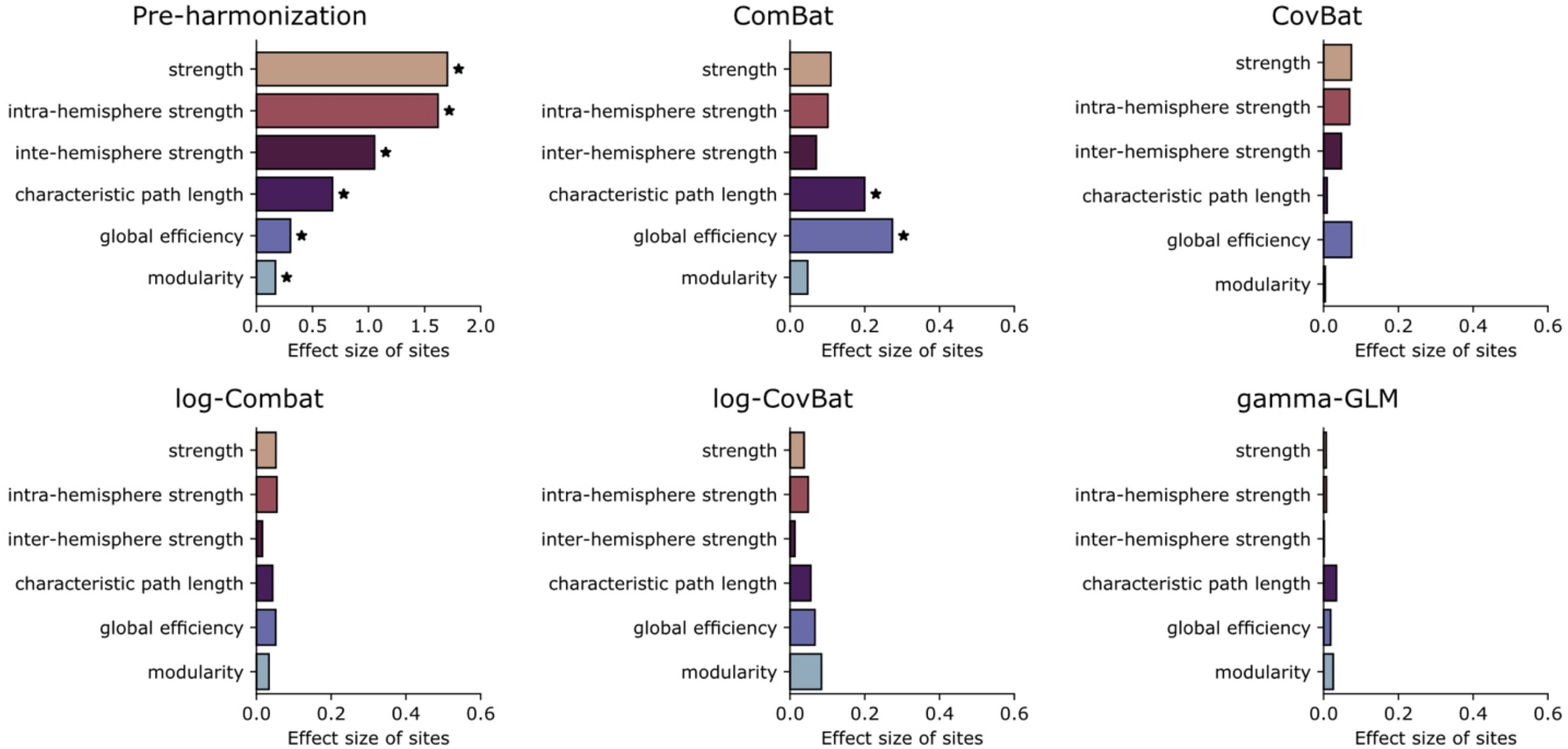
Effect size of site differences on global graph topological measures in the paired PNC-CAR cohort (Data Configuration 1). For each harmonization method, the Cohen’s d effect sizes of sites were evaluated on six global graph topological measures (global strength, intra- and inter-hemisphere strength, characteristic path length, global efficiency, modularity). Significant site effects were indicated by asterisks (p < 0.05, two-sample t-test).

We then explored the site effects on nodewise graph topological features. Figure 5 presents the effect sizes of sites on four node-level graph features, with significant site effects marked by asterisks. The number of nodes with significant site effects was indicated in the top left corner of each plot. While all tested harmonization approaches addressed most site effects on node strength and clustering coefficient, ComBat and CovBat failed to correct site effects on betweenness centrality and local efficiency, leaving a substantial number of nodes with significant site effects (p<0.05, two-sample t-test).

**Figure 5.**
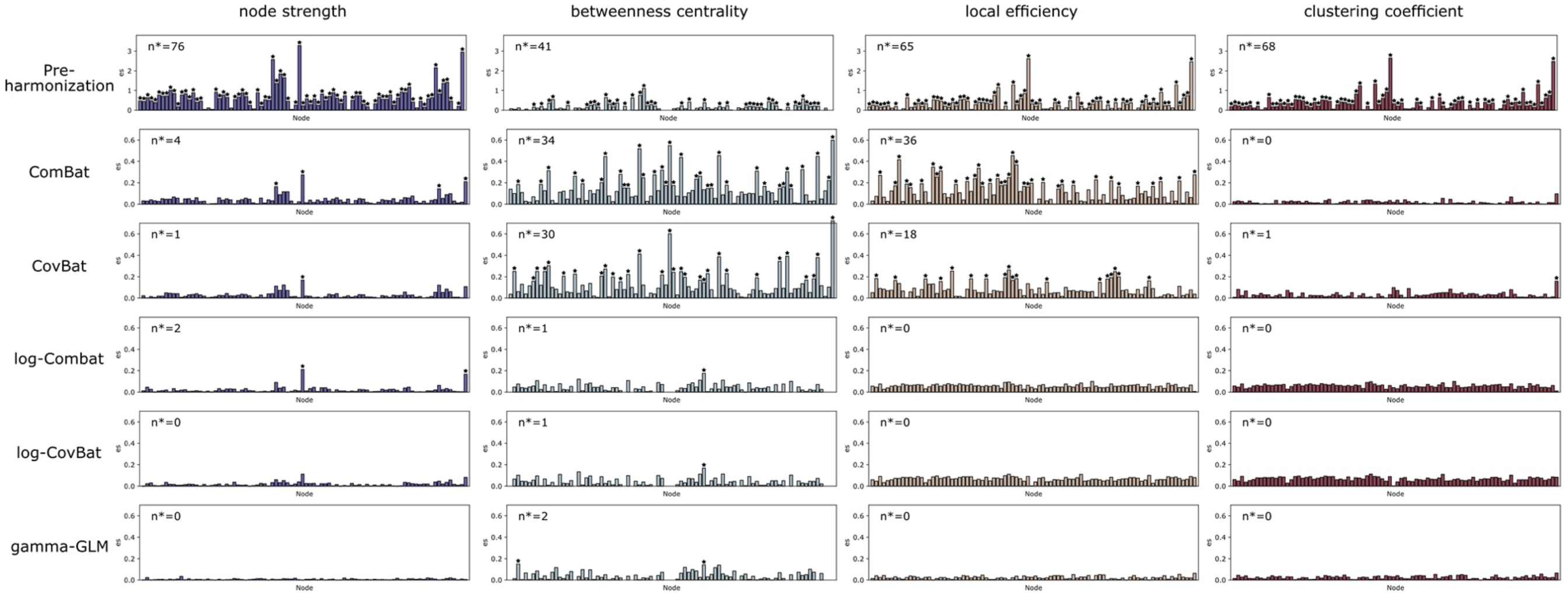
Effect size of site differences on nodal graph topological measures in the paired PNC-CAR cohort (Data Configuration 1). Four nodewise topological measures (node strength, betweenness centrality, local efficiency and clustering coefficient) were evaluated. Significant site effects (p < 0.05, two-sample t-test) were indicated by asterisks. The number of nodes with significant site effects was noted by n* in the top leB corner of each plot.

### 3.4. Evaluation of Biological Variability Preservation

We assessed the replicability of age-related biological variability after applying different harmonization methods. For each site, Spearman correlations between edgewise connectivity strength and subjects’ chronological age were calculated before and after harmonization, controlling for sex as a covariate. Figure 6A shows the edgewise correlations for PNC (first row) and CAR (second row) with paired subjects (Data Configuration 1) before harmonization, displaying only edges with significant age associations (p < 0.05, Spearman R). The differences in edgewise correlation with ages between harmonized and pre-harmonization data were displayed for each harmonization approach. Notably, the gamma-GLM method most closely resembled the pre-harmonization patterns of age associations. Table 4 quantifies the replicability of these age associations using the true positive rate (TPR) of edges that detected significant correlations, compared to the pre-harmonization connectomes. Our results showed that gamma-GLM outperformed other methods by recovering all the positive and negative correlations in the original structural connectomes across both sites, followed by log-ComBat and log-CovBat.

**Table 4.** Recovery of connections with significant age correlation.

| Cohort | Site | # positive correlations | TPR of positive correlations |  |  |  |  | # negative correlations | TPR of negative correlations |  |  |  |  |
| --- | --- | --- | --- | --- | --- | --- | --- | --- | --- | --- | --- | --- | --- |
|  |  | pre-harmonization | ComBat | CovBat | log-ComBat | log-CovBat | gamma-GLM | pre-harmonization | ComBat | CovBat | log-ComBat | log-CovBat | gamma-GLM |
| Paired<br>PNC-CAR | PNC | 245 | 93.47% | 92.65% | 99.59% | 97.14% | 100% | 179 | 79.89% | 79.33% | 97.77% | 97.21% | 100% |
|  | CAR | 223 | 73.54% | 73.54% | 90.13% | 86.55% | 100% | 292 | 88.36% | 89.73% | 94.86% | 94.52% | 100% |
| Confounded<br>NYU-TCD | ABIDEII-NYU | 16 | 75.00% | 81.25% | 87.50% | 93.75% | 100% | 105 | 87.62% | 76.19% | 92.38% | 83.81% | 100% |
|  | ABIDEII-TCD | 35 | 80.00% | 65.71% | 94.29% | 74.29% | 100% | 93 | 82.80% | 67.74% | 94.62% | 91.40% | 100% |

Abbreviations: TPR, true positive rate; # positive (or negative) correlations, number of connections with positive (or negative) age correlations

**Figure 6.**
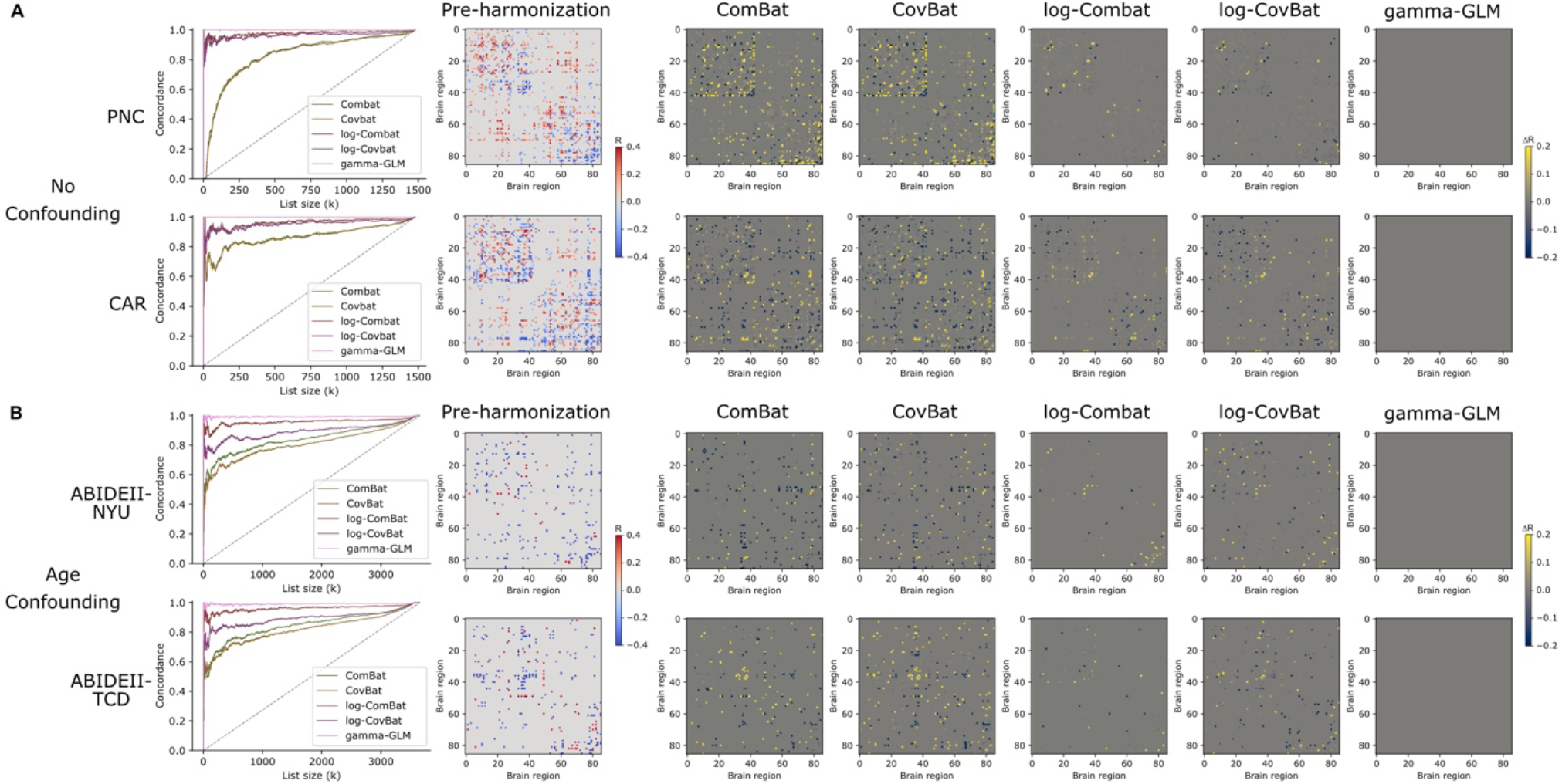
Visualization of edgewise age associations before and aBer applying different harmonization methods. Two scenarios were tested: (A) the paired PNC-CAR cohort (Data Configuration 1) and (B) the confounded NYU-TCD cohort (Data Configuration 2). For each scenario, edgewise correlations between connectivity strength and age were shown for each site before harmonization, displaying only significant age-associated connections (p < 0.05, Spearman R). The differences in edgewise correlations between harmonized data and pre-harmonization data were shown for each harmonization approach. The CAT curves visualized the concordance of edgewise age associations before and aBer harmonization for each method. A CAT curve closer to one indicated better overlap in age associations.

We visualized the replicability of age associations with CAT plots, with each curve representing the concordance of rankings for all edgewise age correlations before and after harmonization, controlling for sex (Figure 6A). In the absence of confounds, gamma-GLM performed the best at both PNC and CAR sites, showing an almost flat CAT curve near 1. The log-ComBat and log-CovBat methods exhibited good overlap, performing behind gamma-GLM, while ComBat and CovBat ranked the last.

### 3.5. Harmonization with Confounding Factors

The previous results were obtained by harmonizing two sites that paired for age and sex to minimize potential confounding of those variables across sites. However, this kind of pairing is not always feasible in multi-site studies, a better harmonization method should be applied on all available data across sites without the pairing process and should be robust to confounding factors. Figure 6B demonstrates the edgewise within-site correlations with age in the confounded NYU-TCD cohort (Data Configuration 2) before and after different harmonization approaches, with ABIDEII-NYU shown in the first row and ABIDEII-TCD in the second row. We noted that, compared to the paired cohort, the confounded cohort demonstrated different edgewise patterns between two sites. Since the two sites included participants from distinctly different age ranges, the age-related effects on structural connectomes may vary. The CAT plots show that the site-specific age-related patterns were best preserved for each site using gamma-GLM harmonization, followed by log-ComBat, log-CovBat, ComBat, and CovBat. The gamma-GLM harmonization successfully recovered all edges with significant positive and negative correlations in presence of confounding factors between age and sites (Table 4).

We calculated the Frobenius distances between the edgewise age correlation patterns of two sites, as shown in Table5. We found the Frobenius distances between two confounded sites were larger than those between two paired sites. Ideally, we expected the between-site distance for age-related patterns should remain the same before and after harmonization. The gamma-GLM most successfully persevered the age-related differences across sites in both paired and confounded scenarios. In contrast, other harmonization approaches inadvertently reduced or increased the age-related differences across sites, likely due to poor fitting caused by an inappropriate choice of distributions for the edges.

Lastly, we evaluated the replicability of age associations when combining two sites. Figure S3 shows the CAT curves for two data configurations. Our results demonstrated that, in the presence of confounding, the replicability of age associations dropped dramatically compared to the paired cohort if sites are combined without proper harmonization, whereas properly harmonized ones maintained high replicability. The gamma-GLM harmonization resulted in highest replicability regardless of age confounds.

### 3.6. Use case 1: Brain age prediction on new datasets

In the following sessions, we evaluated the effectiveness of different harmonization methods in addressing several common challenges that encountered in multi-site structural connectome studies. For this analysis, we used the entire TDC cohort (Data Configuration 3). We first repeated the previous evaluations to examine site effects on edgewise connectivity strength and graph topological measures, both before and after harmonization. The results are presented in Figure S3-S5 in the Supplementary Materials.

We trained a brain age predictor using an SVR model on the training set of PNC and apply it to the remaining sites before and after different harmonization methods. Table 6 shows the model’s prediction performance at each new site, as well as for the combined data from all new sites. For reference, the performance of the trained brain age predictor on the testing set of PNC was provided (underlined). Our results demonstrated that using proper harmonization to remove site-specific variations between PNC and new sites could largely improve the generalizability of the brain age predictor to new datasets. The gamma-GLM-harmonized data achieved the lowest MAE and RMSE, along with the highest Pearson R correlation, across most new sites with unseen data. Although log-ComBat attained the lowest MAE and RMSE in two sites (Lurie MEGlab, ABIDEII-TCD), it did not match gamma-GLM’s performance in terms of Pearson R correlation.

**Table 5.** Frobenius distances between site-specific edgewise patterns of age associations.

| Pairwise Site differences | Pre-harmonization | ComBat | CovBat | log-ComBat | log-CovBat | gamma-GLM |
| --- | --- | --- | --- | --- | --- | --- |
| Paired<br>PNC-CAR | <u>14.4747</u> | 16.0879<br>(+1.6132) | 16.1088<br>(+1.6341) | 14.3714<br>(-0.1032) | 14.3696<br>(-0.1051) | **14.4743**<br>**(-0.0004)** |
| Confounded<br>NYU-TCD | <u>19.4563</u> | 20.6006<br>(+1.1443) | 20.2229<br>(+0.7666) | 19.3874<br>(-0.0689) | 19.5346<br>(+0.0783) | **19.4563**<br>**(-0.0000)** |

**Table 6.** Impact of harmonization on the generalizability of brain age predictors to new datasets.

| Dataset | Metric | Pre-harmonization | ComBat | CovBat | log-ComBat | log-CovBat | gamma-GLM |
| --- | --- | --- | --- | --- | --- | --- | --- |
| PNC | MAE | <u>1.7321</u> | <u>1.6752</u> | <u>1.6433</u> | <u>1.7078</u> | <u>1.7001</u> | <u>1.7281</u> |
|  | RMSE | <u>2.1681</u> | <u>2.0683</u> | <u>2.0400</u> | <u>2.1396</u> | <u>2.1329</u> | <u>2.1651</u> |
|  | Pearson R | <u>0.7405</u> | <u>0.7662</u> | <u>0.7734</u> | <u>0.7483</u> | <u>0.7504</u> | <u>0.7414</u> |
| CAR | MAE | 3.7216 | 2.4807 | 2.4579 | 1.9644 | 1.9739 | **1.8536** |
|  | RMSE | 4.3141 | 3.1131 | 3.0585 | 2.3928 | 2.4044 | **2.2631** |
|  | Pearson R | 0.0548 | 0.3260 | 0.3175 | 0.6400 | 0.6446 | **0.7074** |
| Lurie MEGlab | MAE | 5.0619 | 2.7072 | 2.5526 | **1.9518** | 2.0532 | 2.0260 |
|  | RMSE | 5.4438 | 3.2332 | 3.0941 | **2.2854** | 2.4712 | 2.3330 |
|  | Pearson R | 0.2194 | 0.2418 | 0.2436 | 0.5063 | 0.4355 | **0.5426** |
| ABIDEII-NYU | MAE | 5.5198 | 3.1685 | 3.1896 | 1.8013 | 2.0526 | **1.7870** |
|  | RMSE | 5.7817 | 3.7338 | 3.9420 | 2.2315 | 2.5171 | **2.2105** |
|  | Pearson R | -0.5111 | -0.2693 | -0.3182 | 0.4385 | 0.3965 | **0.4676** |
| ABIDEII-SDSU | MAE | 2.8504 | 3.4530 | 3.4925 | 2.5960 | 2.6739 | **2.3822** |
|  | RMSE | 3.3662 | 4.1185 | 4.1908 | 3.1410 | 3.2959 | **2.9947** |
|  | Pearson R | 0.2313 | -0.4330 | -0.4747 | 0.2078 | 0.1278 | **0.3514** |
| ABIDEII-TCD | MAE | 2.5921 | 2.5443 | 2.9942 | **2.0126** | 2.1363 | 2.0621 |
|  | RMSE | 3.0745 | 3.2309 | 3.6377 | **2.3909** | 2.5667 | 2.4667 |
|  | Pearson R | 0.5522 | 0.0682 | -0.1160 | 0.6647 | 0.5909 | **0.6804** |
| All New Sites | MAE | 3.9080 | 2.6690 | 2.6702 | 2.0137 | 2.0719 | **1.9419** |
|  | RMSE | 4.4883 | 3.3017 | 3.3100 | 2.4460 | 2.5346 | **2.3653** |
|  | Pearson R | 0.2027 | 0.3935 | 0.3433 | 0.6790 | 0.6587 | **0.7201** |

<sup>1</sup>The performance of brain age predictors is reported on the testing set of PNC (underlined). The brain age predictors are directly applied to other sites to evaluate the generalizability of the model to new datasets.
<sup>2</sup>The best-performing harmonization method for each metric is highlighted in bold.
<sup>3</sup>Abbreviations: MAE, mean absolute error; RMSE, root mean square error; Pearson R, Pearson's correlation.

### 3.7. Use case 2: Enhancing statistical power for ASD studies

In use case 2, we evaluated the effectiveness of harmonization methods in the pooled multi-site ASD cohort (Data Configuration 4) to examine their impact on detecting clinical group differences. Our findings indicated that prior to harmonization, the structural connectome data showed no significant group-level differences between ASD and TDC groups, nor any group-by-age interactions in global graph measures (p>0.05, FDR-adjusted). Additionally, none of the global graph measures demonstrated significant associations with clinical scores (p>0.05, FDR-adjusted). Post harmonization, we found that males with ASD showed significantly higher intra-hemispheric strength compared to the TDC males (Cohen’s d=0.37, p=0.041, FDR-adjusted). Significant group-by-age interactions were observed for clustering coefficient (standardized ***β***=0.27, p=0.005, FDR-adjusted) and global efficiency (standardized ***β***=0.26, p=0.006, FDR-adjusted), with both measures increasing with age in ASD but decreasing in TDC. Intra-hemispheric strength showed a marginally significant correlation with the SRS-2 T-score (R=0.12, p=0.072, FDR-adjusted), while clustering coefficient (R=0.15, p=0.038, FDR-adjusted) and global efficiency (R=0.14, p=0.038, FDR-adjusted) were significantly and positively correlated with the VABS communication subscale. However, ADOS and the two other domains of VABS (socialization and daily living) did not correlate with any global graph measures (p>0.05, FDR-adjusted).

## 4. Discussion

The scientific community is transitioning into the era of big data, where multiple research institutions and clinical centers collaborate to collect and maintain large-scale datasets. This study tackles the critical challenge of effectively merging multi-site structural connectome data without introducing site-related biases. Our harmonization framework provides a way to expand sample sizes, enabling a comprehensive representation of both healthy controls and patient populations to support more robust findings in the field. While previous studies introduced harmonization approaches at the level of dMRI signals [23, 26], they did not remove site effects at the level of the structural connectome [27], introducing bias in downstream graph analyses and other connectome-based investigations. Our study addressed the harmonization of the connectomes.

We provide practical guidance for researchers aiming to harmonize structural connectome data, following the pipeline outlined in Figure 1. Overall, we recommend the gamma-GLM approach as our top choice, as it outperformed other models in removing site effects while effectively preserving both demographic and disorder-related biological integrity throughout the harmonization process. In contrast, we found that harmonization techniques based on normal distributional assumption failed to remove site effects consistently for all connections in connectome data, potentially resulting in spurious findings in downstream analyses. Given the unique distributional properties of connectivity strength, it is essential to validate the underlying data assumptions when applying harmonization methods. Standard harmonization methods in neuroimaging, such as ComBat, CovBat, or simple linear regression, assume a normal distribution and are therefore unsuitable for structural connectomes, where most edges follow a gamma distribution. Moreover, ComBat and CovBat can yield negative values after harmonization, which lack physical meaning in structural connectomes, where edges represent (normalized) numbers of streamlines connecting brain regions. Applying these methods without careful consideration and validation [35] could lead to incorrect conclusions or unintended consequences in multi-site studies.

Notably, connectomes represent graph-structured brain connectivity, so harmonization should not only remove edgewise site effects but also eliminate site-related variations in their graph properties, commonly measured by graph topological measures [57]. Most prior work only evaluated graph measures at the global level [31, 35], but our results indicated that even though no site differences were observed in global measures, site differences can still exist at the nodal level. For example, while CovBat effectively removed site effects in all common global graph measures, substantial site effects persisted in nodal measures such as betweenness centrality and local efficiency. Therefore, we recommend evaluating harmonization effectiveness for structural connectomes across all three levels—edgewise connectivity values, nodal, and global graph measures—to ensure that site effects are adequately eliminated prior to downstream analyses. Our results showed that the harmonization framework based on gamma-GLM performed best across all three levels, followed by log-CovBat and log-ComBat approaches with our adaptation to structural connectomes.

The primary goal of harmonization is to integrate data across sites to facilitate the investigation of biologically based hypotheses using larger and more comparable structural connectome data. Therefore, it is essential that biologically relevant properties are not inadvertently altered or removed during the harmonization process. Given that age-related connectivity changes are critical in many studies of neurodevelopment and neurodegenerative processes [62, 69], we evaluated edgewise associations with age, both pre- and post-harmonization. Our findings indicated that gamma-GLM-based harmonization framework most effectively preserved these biologically meaningful associations across sites and remained robust even when age was confounded with site effects. The gamma-GLM model provides a better fit for covariate effects in structural connectome data, further emphasizing its advantage over other approaches for multi-site connectome data.

We demonstrated the practical utility of structural connectome harmonization with two use cases. The first use case addressed a commonly reported challenge in collaborative research: predictive models developed with one dataset often fail to predict unseen data from new sites [70, 71]. This occurs because new datasets may exhibit substantial differences in scanners, acquisition protocols, and other site-related effects compared to the training cohort, limiting the generalizability of predictive models across sites. Here, we created machine learning models to predict structural connectome-based brain age using data from one site and evaluate the generalizability of these predictors on other sites. We showed that harmonizing structural connectomes across sites, particularly using the gamma-GLM based model, enhanced the generalizability of predictive models to unseen data at new sites. This approach can be further applied to a federated learning setup in healthcare [72, 73]. In this context, researchers can first harmonize and standardize decentralized data sources to a central site to ensure comparability, thereby improving further generalizability when deployed from and to the central site.

The second use case of harmonization is to increase the statistical power of case-control cohort studies via pooling structural connectomes from multiple sources. This approach is especially beneficial for clinical studies involving hard-to-recruit clinical populations, where data from a single site is often insufficient. By pooling patient data from various sources, researchers can create larger and more diverse cohorts, which are crucial for detecting subtle connectivity differences associated with various clinical conditions. To achieve this, the effectiveness and robustness of harmonization methods must be carefully evaluated within the context of the patient population. Our results, based on a clinical cohort of ASD participants and controls, demonstrated that without proper harmonization, site-related effects could obscure associations between graph topological measures and neurodivergent traits. While there is no definitive ground truth for *accuracy* in such measurements, harmonization allowed sufficient improvement in estimation *precision* by mitigating noise introduced by site and scanner differences. In the present data, harmonization with gamma-GLM effectively resolved significant differences in structural connectomes between diagnostic groups that were previously undetectable.

Despite such a comprehensive set of experiments evaluating the pros and cons of different frameworks for connectome harmonization, there are several issues that warrant attention in future research. Users should note that the gamma-GLM approach does not model site-specific variance and covariance. When image acquisition and reconstruction protocols vary significantly across sites, careful checks are necessary to ensure that such variations do not affect the data properties [55]. Although we did not observe significant site-related differences in covariance in this study, these effects may emerge in subnetwork analyses or when structural connectomes are modulated by diffusion metrics like fractional anisotropy (FA) or mean diffusivity (MD). While we tackled the most used definition of structural connectomes and tested common distributions from the exponential family, researchers creating connectomes using more complex methods may need to explore additional distributions (e.g., zero-adjusted gamma distribution) and other statistical metrics. In such cases, more flexible distributional regression models, like the Generalized Additive Model for Location, Scale, and Shape (GAMLSS) [74], might be suitable. Another area for future work lies in expanding demographic variables involved during harmonization. Incorporating factors like ethnicity and socioeconomic status could enhance the applicability of findings across diverse populations to promote health equity [75, 76]. Our harmonization and evaluation framework can also be adapted to better address the specific goals of structural connectome studies in these contexts.

## 5. Conclusion

In summary, we outlined a path forward for harmonizing multi-site structural connectome datasets, thereby facilitating collaborative clinical research. Given the complex nature of connectomes, it is essential to examine the distributional properties of the connectome data. The appropriate harmonization framework, such as gamma-GLM or more flexible distributional regression models, should be selected to ensure that the integrity of the data is maintained post-harmonization. Additionally, all harmonization results should be carefully assessed for site effects at different levels of measures, such as edgewise connectivity strength and global and nodal-level graph properties, to ensure an unbiased interpretation of the data in downstream connectome studies.

## Supporting information

Supplementary Results

## Notes

### Competing Interest Statement

The authors declare the following potential conflicts of interest:
Timothy P.L. Roberts holds stock in Prism Clinical Imaging, has a partnership interest in Proteus Neurodynamics, and has received consulting fees from Fieldline Inc. and WestCan Proton Therapy.
Russell T. Shinohara has received consulting fees from Octave Bioscience and the American Medical Association.
All other authors (Rui S. Shen, Drew Parker, Andrew A. Chen, Birkan Tunc, Benjamin E. Yerys, and Ragini Verma) report no conflicts of interest.

### Summary of Updates

add supplementary materials; update the title; improve the writing

