## Supplementary Results for "Big Data, Small Bias: Harmonizing Diffusion MRI-Based Structural Connectomes to Mitigate Site-Related Bias in Data Integration"

**Supplementary Material**

---

### 1. Supplementary Results

#### *1.1. Structural connectome distributions in five additional sites*

We evaluated the distributional properties of structural connectomes for each diagnostic group (ASD and TDC) within each dataset. Each edge of the pre-harmonized structural connectomes was assessed to determine whether it conformed to one of three hypothesized probability distributions: normal, log-normal, or gamma. Our analysis revealed that the gamma distribution provided the best fit across diagnostic groups and sites, exhibiting significantly lower KS distances compared to the other two distributions ($p < 0.0001$, paired t-test), with the log-normal distribution ranking second.

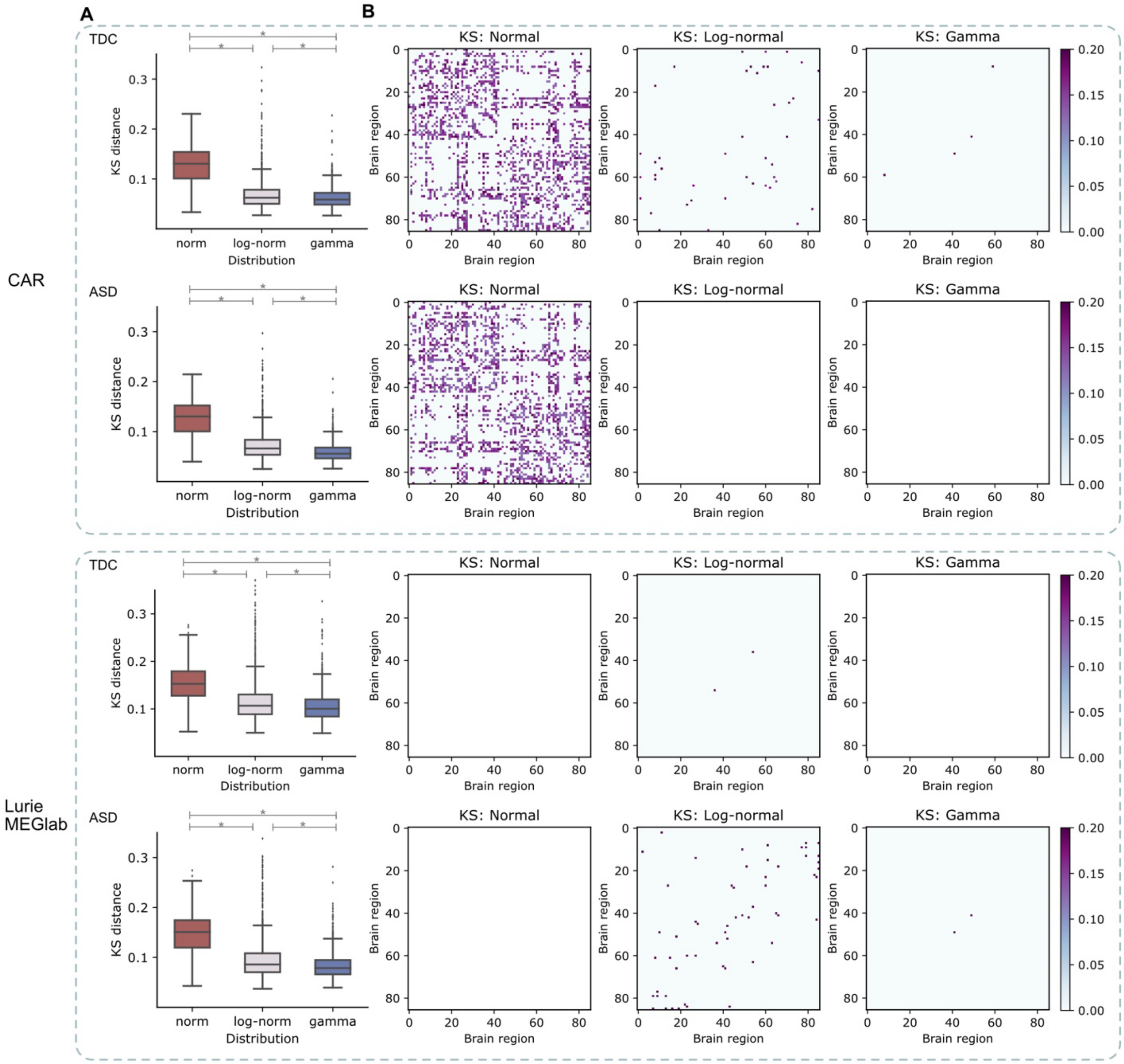


*Figure S1 Goodness of fit test for each dataset. A) Edgewise KS distances between the observed and three hypothesized distributions (normal, log-normal, and gamma) of connectivity strength for each diagnostic group in each dataset. Asterisks*

*denote significant differences between those hypothesized distributions (p < 0.0001, paired t-test). The gamma distribution provided a significantly better fit than the other two hypothesized models. B) Heatmaps of edgewise KS distances, showing only the edges with significant discrepancy from the observed distributions in pre-harmonized structural connectomes (p < 0.05, FDR-adjusted).*

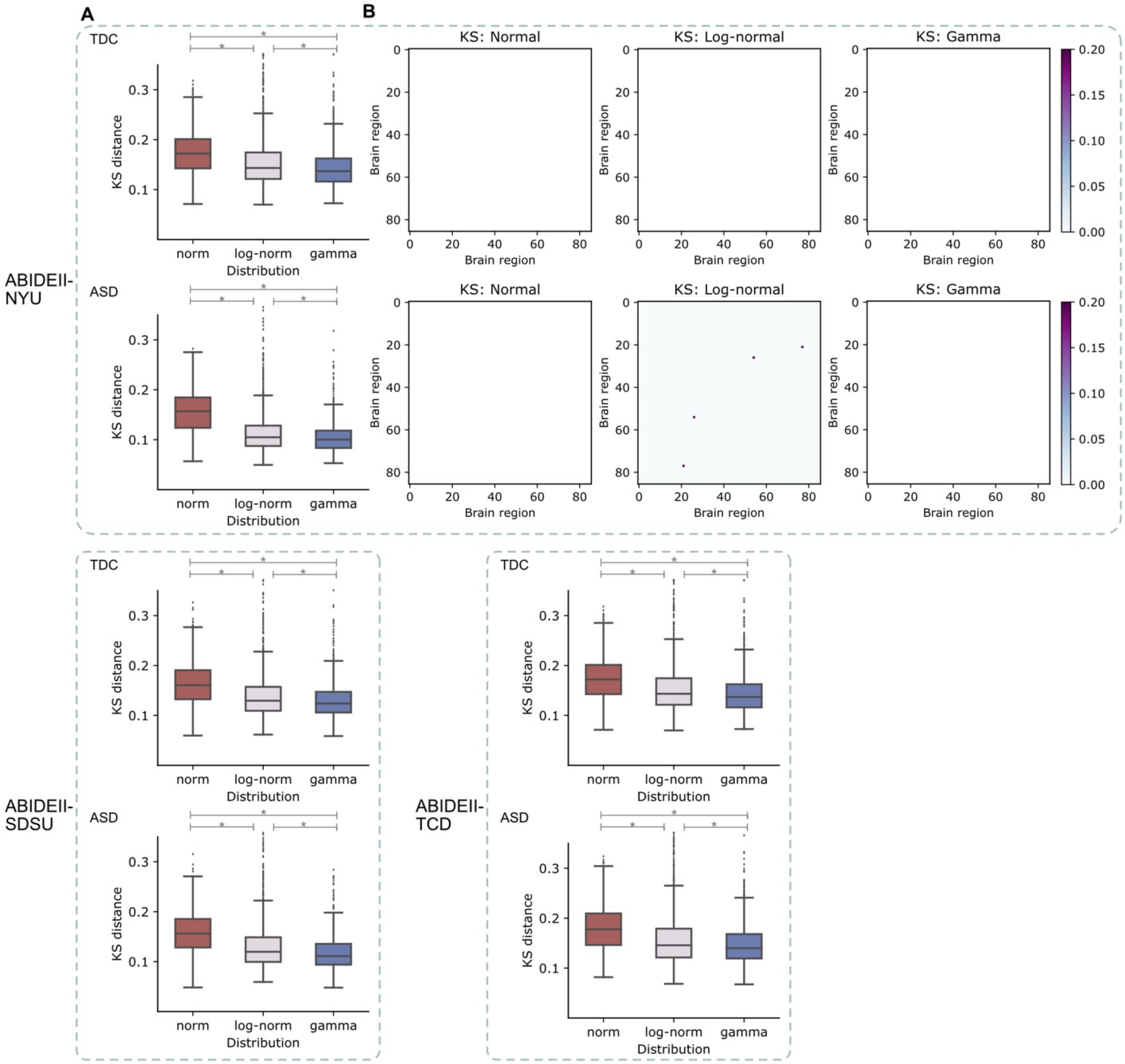


*Figure S1 (continued) Goodness of fit test for each dataset.*

Heatmaps of KS distances were also generated for each edge, highlighting only those edges with significant deviations from the observed distributions (p < 0.05, FDR-adjusted). In the CAR dataset, the normality assumptions underlying methods like ComBat and CovBat were violated for most edges, as the structural connectome data more closely followed a gamma distribution. At the Lurie MEGlab and three ABIDEII sites, the small sample sizes limited the statistical power of the edgewise Kolmogorov–Smirnov tests, leading to most edges failing to pass multiple comparison corrections for significant deviations from the hypothesized distributions.

*1.2. Analysis of different consistency levels*

In the primary analyses, we applied a consistency-based thresholding at a density level of 40% to filter out spurious connections after probabilistic tractography, retaining only the most consistent connections across the population. Here, we evaluated the distributional properties of edges by varying consistency levels (Top 10%, 10%-20%, ..., 80%-90%). For each analysis, edges within each consistency-level bin were selected, and the goodness of fit of three hypothesized probability distributions was examined. While all sites were included in this analysis, the TDC and ASD groups were evaluated separately. The primary analyses corresponded to retaining edges from the top four bins (Top 10%, 10%-20%, 20%-30%, and 30%-40%).

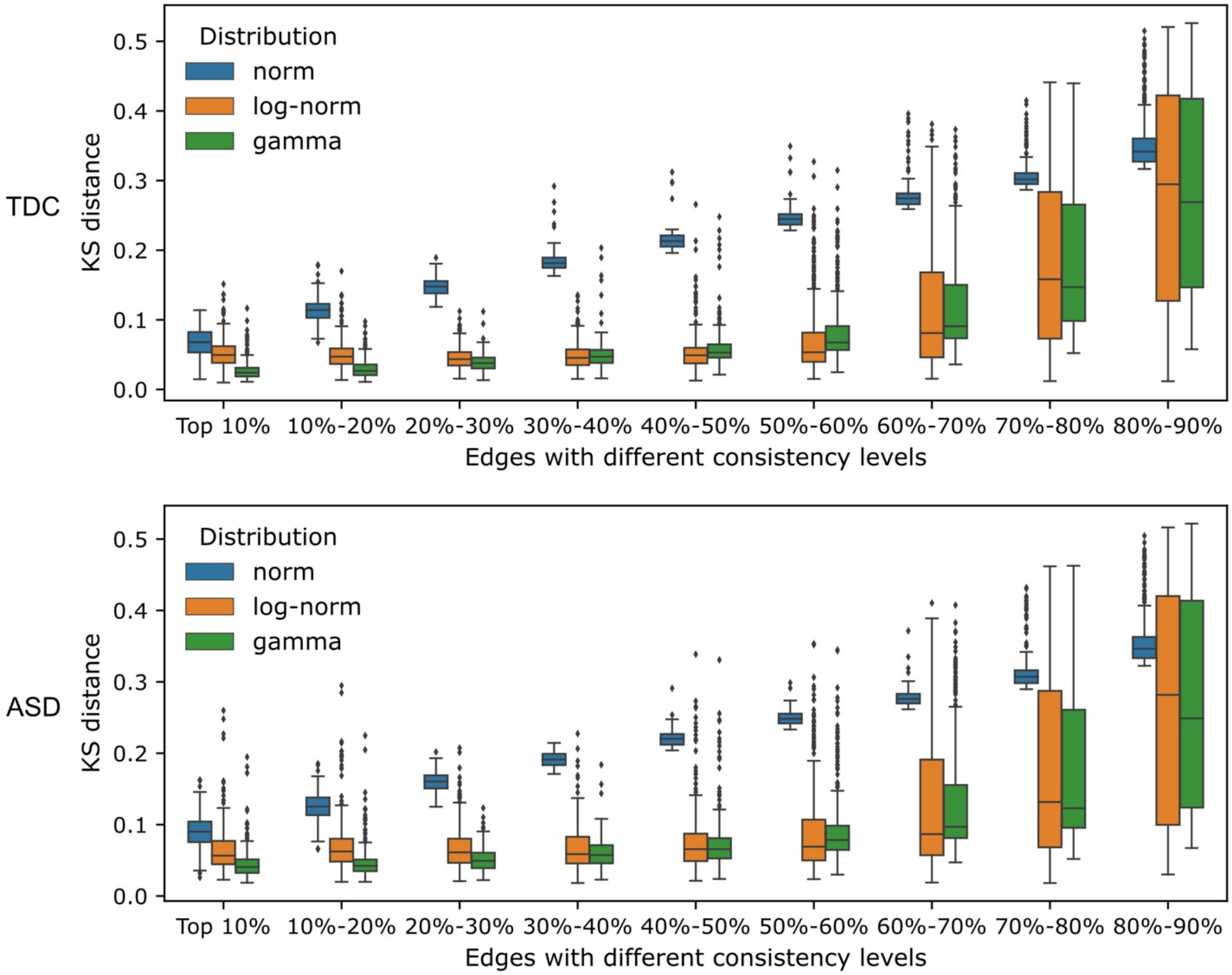


*Figure S2 Goodness of fit test for edges with different consistency levels. Edges from each consistency-level bin were selected, and the KS distances to three hypothesized distributions were calculated. The primary analyses (40% density) corresponded to retaining edges from the top four bins (Top 10%, 10%-20%, 20%-30%, and 30%-40%).*

*1.3. Joint assessment of replicability of age associations after combining sites*

In the primary analyses, we assessed the replicability of edgewise age associations separately within each site. In this analysis, we tested the replicability of age associations by combining data from two sites after harmonization. We considered two cases: combining paired PNC-CAR data in the absence of age confounds (Data Configuration 1) and combining age-confounded NYU and TCD data (Data Configuration 2). Additionally, we included a scenario where the two sites were merged without harmonization to examine the effect of harmonization in addressing confounding factors during data integration.

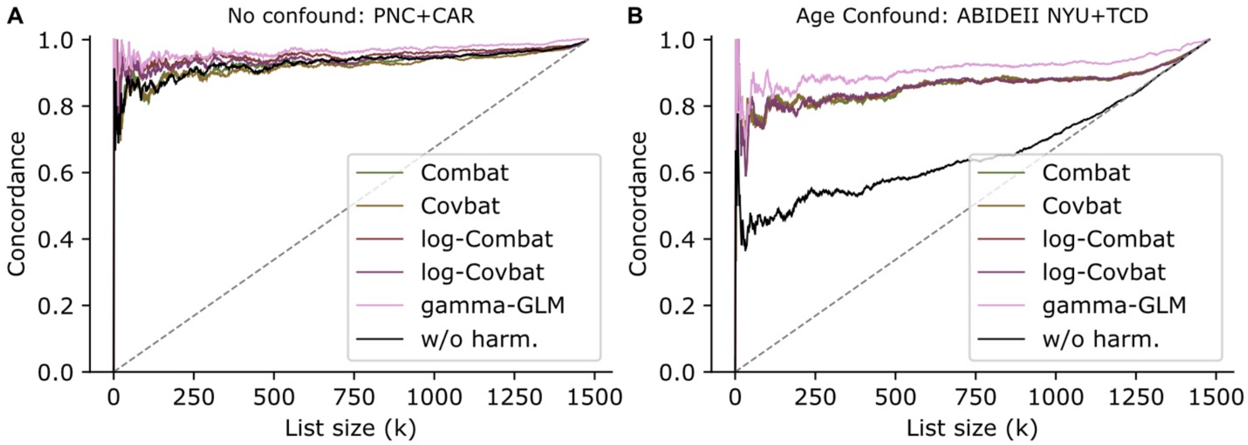


*Figure S3 CAT curves illustrating the joint assessment of replicability of age associations after combining sites. Data configurations include A) paired PNC-CAR data in the absence of age confounds (Data Configuration 1), and B) age-confounded NYU and TCD data (Data Configuration 2). We also tested a scenario where the two sites were combined without harmonization (w/o harm.). In the presence of confounding, the replicability of age associations significantly decreases when sites are merged without harmonization, whereas harmonized data maintain high replicability. The gamma-GLM harmonization resulted in highest replicability regardless of age confounds.*

#### *1.4. Evaluation of harmonization on the entire TDC cohort*

We repeated the evaluation of harmonization approaches on the entire TDC cohort at three levels: edgewise connectivity values, nodal measures, and global graph metrics. Overall, gamma-GLM outperformed other methods across all levels.

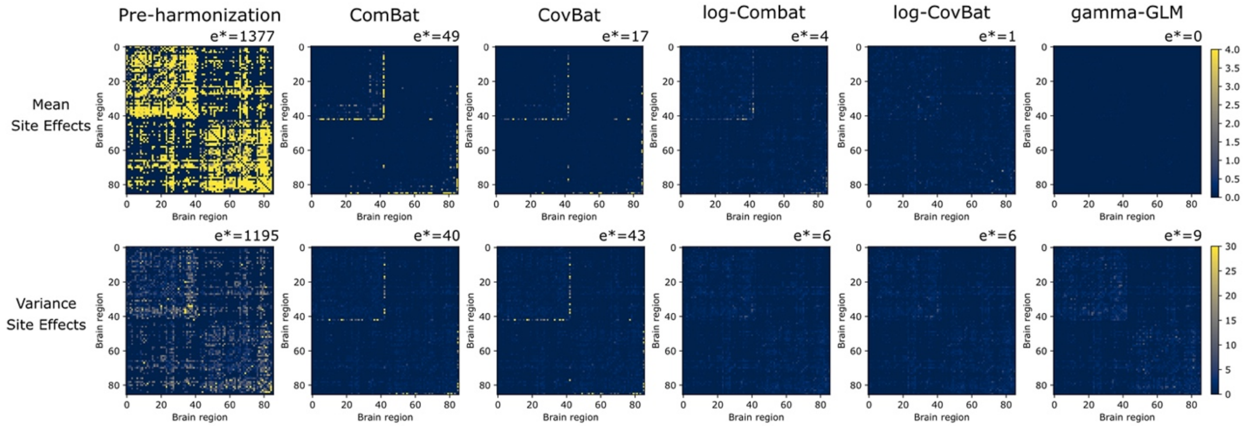


*Figure S3 Evaluation of edgewise site effects on mean (first row) and variance (second row) connectivity strength using the entire TDC cohort (Data Configuration 3), showing the Kruskal-Wallis H statistics for mean site effects and F* statistics from the Brown-Forsythe test for variance site effects. The number of edges with significant site effects was noted by e* in the top left corner of each plot.*

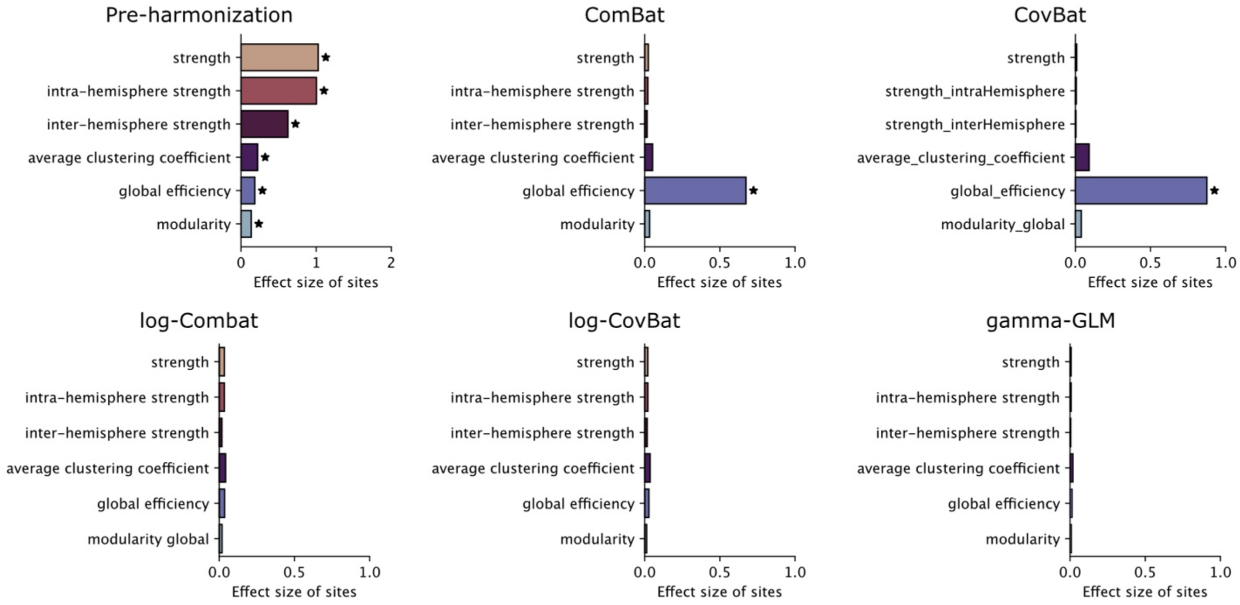


*Figure S4 Effect size of site differences on global graph topological measures in the entire TDC cohort (Data Configuration 3). For each harmonization method, the Cohen's f effect sizes of sites were evaluated on six global graph topological measures. Significant site effects were indicated by asterisks ($p < 0.05$, one-way ANOVA test).*

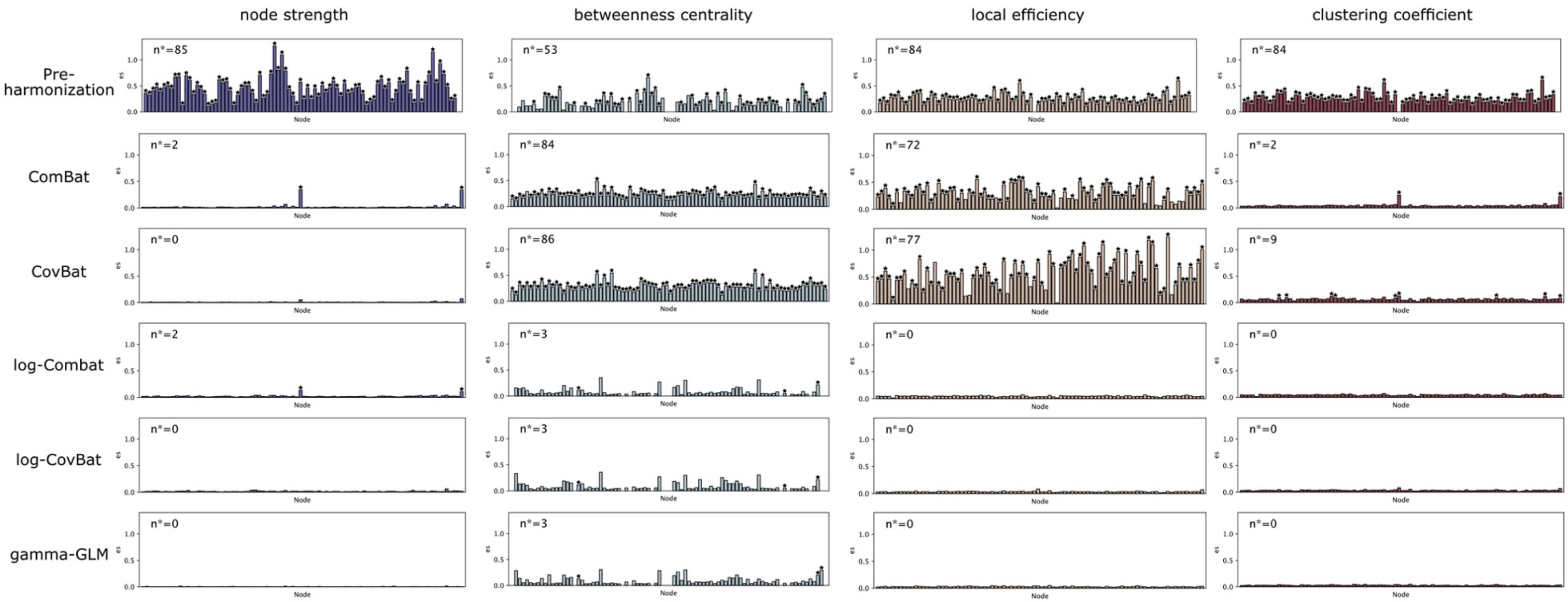


*Figure S5 Effect size of site differences on global graph topological measures in the entire TDC cohort (Data Configuration 3). For each harmonization method, the Cohen's f effect sizes of sites were evaluated on six global graph topological measures. Significant site effects were indicated by asterisks (p < 0.05, one-way ANOVA test).*
